# Personally tailored cancer management based on knowledge banks of genomic and clinical data

**DOI:** 10.1101/057497

**Authors:** Moritz Gerstung, Elli Papaemmanuil, Inigo Martincorena, Lars Bullinger, Verena I Gaidzik, Peter Paschka, Michael Heuser, Felicitas Thol, Niccolo Bolli, Peter Ganly, Arnold Ganser, Ultan McDermott, Konstanze Döhner, Richard F Schlenk, Hartmut Döhner, Peter J Campbell

## Abstract

Sequencing of cancer genomes, or parts thereof, has become widespread and will soon be implemented as part of routine clinical diagnostics. However the clinical ramifications of this have not been fully assessed. Here we assess the utility of sequencing large and clinically well-annotated cancer cohorts to derive personalized predictions about treatment outcome. To this end we study a cohort of 1,540 patients with AML (acute myeloid leukemia) with genetic profiles from 111 cancer genes, cytogenetic data and diagnostic blood counts. We test existing and develop new models to compute the probability of six different clinical outcomes based on more than 100 genetic and clinical variables. The predictions derived from our knowledge bank are more detailed and outperform strata currently used in clinical practice (concordance C=72% v C=64%), and are validated on three cohorts and data from TCGA (C=70%). Our prognostic algorithm is available as an online tool (http://cancer.sanger.ac.uk/aml-multistage). A simulation of different treatment scenarios indicates that a refined risk stratification could reduce the number of bonemarrow transplants by up to 25%, while achieving the same survival. Power calculation show that the inclusion of further genes most likely has small effects on the prognostic accuracy; increasing the number of cases will further reduce the error of personalized predictions.

---

Led by a small number of high-profile successes, there has been considerable enthusiasm for the concept of personally tailoring cancer management based on individual genomic profiles^1,2^. Mutations in cancer genes fundamentally drive the tumor’s growth; applications of genomics in cancer medicine include enhanced diagnostic accuracy through molecular characterization, personalized forecasts of a given patient’s prognosis and support for choosing between different therapeutic options ^3,4^. There are complications to this narrative. Surprisingly few cancer genes are straightforward therapeutic targets. Many cancer genes are only rarely mutated in a given tumor type. Each patient’s tumor typically has several driver mutations. Above all other complications, though, is the challenge that there are hundreds to thousands of different combinations of driver mutations observed across patients for most tumor types ^5–7^.

The promise of precision medicine has triggered considerable funding commitments, such as the Precision Medicine Initiative in USA, Genomics England in UK and similar efforts in several other countries ^8,9^. Central to these initiatives is the concept of a so-called “cancer knowledge network” ^9^, in which molecular data on patients’ cancers will be matched with their clinical outcomes to enable personalized management strategies. Despite these investments reaching hundreds of millions of dollars in scale, there has been little formal evaluation of the potential utility of knowledge banks. In particular, it is unclear whether accurate predictions about cancer outcomes can be made from a large genomic-clinical database; what improvements in survival at the population level might be achieved from personally tailored therapeutic choices; and what sample sizes knowledge banks need to accrue before predictions are sufficiently accurate to underpin decision support for the individual patient.

Here, we explore these questions using genomic and clinical data from 1,540 patients with acute myeloid leukemia (AML) undergoing intensive treatment. This exemplar is interesting because of a real, current therapeutic dilemma for these patients – who should be offered an allogeneic hematopoietic cell transplant (allograft) in first complete remission (CR1) ^10,11^? The equations are not straightforward. Allogeneic hematopoietic cell transplants in first complete remission undoubtedly decrease relapse rates for most patients, but this comes at the cost of higher treatment-related mortality, as high as 20-25% at 3 months ^12^, with a further 30% risk of debilitating chronic morbidity ^13^. Furthermore, even though more patients relapse after chemotherapy in first remission, up to a fifth can then be successfully salvaged with allografts or more intensive chemotherapy ^14,15^.

In the following we present a formal evaluation of a per-patient risk assessment demonstrating that overall survival can be predicted twice as accurately compared to current risk stratification schemes. We devise a multistage prognostic framework capable of quantifying the odds of multiple concurrent clinical endpoints, such as treatment associated mortality and relapse rates. Applied to the dilemma of stem cell transplants, this analysis indicates that potentially 1/4 of transplants could be saved using precision medicine approaches.

## RESULTS

### Predicting complex patient outcomes from genomic and clinical variables

We recently studied 1540 patients with AML from three clinical trials of intensive treatment run by the German-Austrian AML Study Group. As described in Ref. 16, we sequenced all coding exons of 111 myeloid cancer genes in diagnostic leukemia samples, identifying driver point mutations and combining these data with the clinical trials database to generate a comprehensive knowledge bank. Here, we focus on evaluating the utility of the knowledge bank for generating predictions about outcomes personally tailored to the individual patient, and how this can be used to compare likelihoods of favorable outcome under different treatment strategies as a route to clinical decision support. All data and analysis code is documented in Supplementary Data 1 and available at github.com/mg14/AML-multistage.

Throughout, we use overall survival as the primary end-point of these analyses since the aim of intensive therapy in the young AML patient is cure. The full dataset consists of 231 predictor variables, spanning the seven broad categories of fusion genes, copy number alterations, point mutations, gene-gene interactions, demographic features, clinical risk factors and treatment received, across 1540 patients. To assess the accuracy of our predictions we use the following validation strategies: (1) random cross-validation on this dataset, (2) building models from any two clinical trials here and testing on the third; and (3) testing the model built from all three German-Austrian trials on an independent AML cohort from USA (TCGA) ^17^.

We tested a range of regularized regression methods for predicting survival and also implemented more novel approaches, based on random effects and multistage statistical models, for deriving detailed associations between genomic and clinical endpoints (Figure 1a; Supplementary Methods). Using a variety of accuracy measures, the random effects models and multistage models typically scored best in predicting overall survival, roughly doubling the R^2^ compared to current strata (Figure 1b-c). A key aspect of these approaches is that all variables are used to generate patient predictions, whereas conventional methods tended to choose reduced subsets of 10-20 variables. These data therefore suggest that a large share of the between-patient variability in prognosis can be captured by the few dominant effects selected in reductionist models, but that there is also a measurable influence on survival from the long tail of infrequently mutated genes and gene-gene interactions.

**Figure 1.**
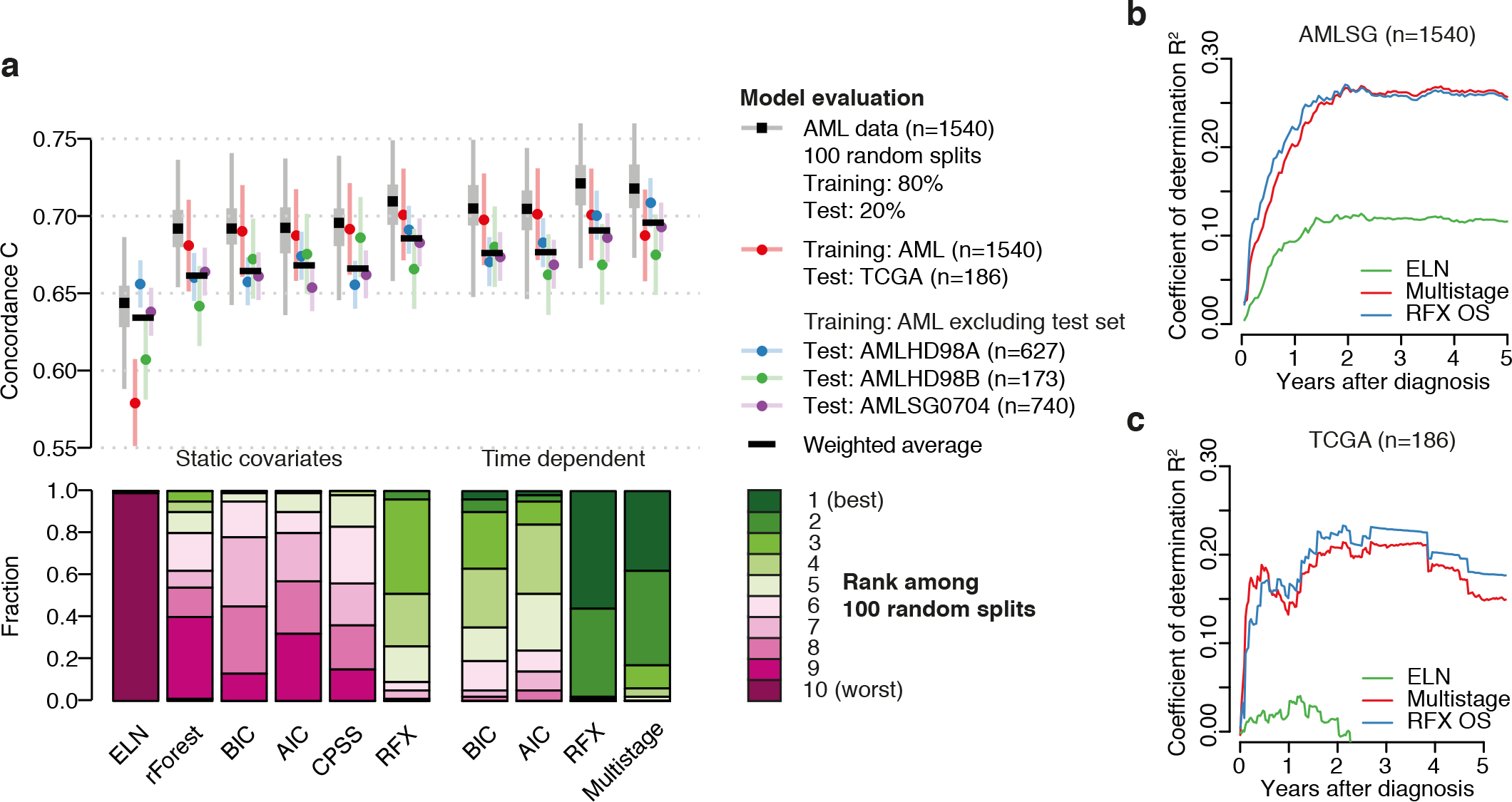
Systematic model comparison. a. Top panel: concordance of different model predictions for overall survival. Shown are distributions across 100 random cross validation splits (grey), as well as point estimates with error bars for predictions evaluated on a single test set (red, blue, green and purple). Predictions for the multistage model are at 3yrs after diagnosis. The lower panel shows the distribution of ranks across the 100 random cross validation splits. b. Coefficient of determination *R*_2_ for time-dependent leave-one-out random effects and multistage predictions of the AMLSG cohort, evaluated at each time (x-axis). c. Same as **b**, evaluated on TCGA data

The multistage model offers the advantage of separating the sources of mortality - death without complete remission, non-relapse death (mostly treatment-related) and death after relapse – and similarly computes the probabilities of survival in induction, first remission (CR1) and after relapse (Figure 2a-b). Briefly, our model estimates the fold change in the rates of 5 possible transitions between the 6 states, based a random effects model, and then computes the time-dependent probability for each patient to be in a given state at a given time (Figure 1c, Supplementary Methods). The analysis shows that the added detail does not come at the cost of overfitting: The combined prediction of overall survival, based on the sum of the all three sources of mortality yields the same accuracy as predicting overall survival using a single transition.

**Figure 2.**
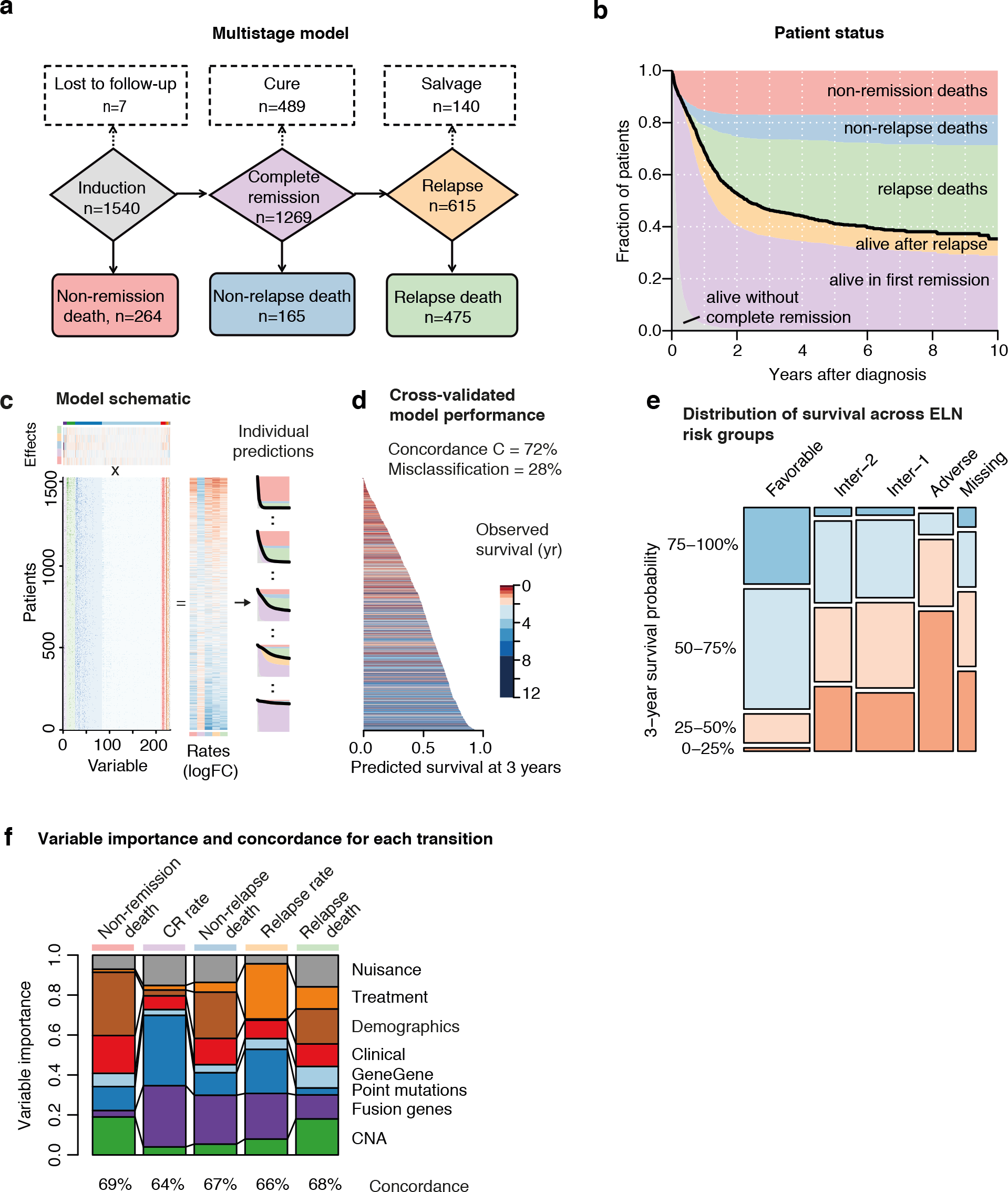
Multi-stage modeling of patient fate. a. Multi-stage model of patient tra ectories. Boxes indicate different stages during treatment, with possible transitions indicated by solid arrows. The model calculates the rate of each transition, based on each patient’s genomic and clinical data. From these rates, the model subsequently derives the probabilities for a patient to be in a particular stage at a given time. Numbers in each box indicate the total number of patients that have entered a given stage in during follow-up. Patient that have not progressed further are shown in the dotted boxes on the top. b. Sediment plot showing the fraction of patients in a given stage at a given time after diagnosis. The thick black line denotes overall survival, which is the sum of the deaths without complete remission (red), non-relapse mortality (blue) and mortality after relapse (green). c. Schematic overview of multistage regression. The model estimates the additive effect of each of 231 prognostic variables on the relative rate of all 5 possible time-dependent transitions shown in (a). The resulting log-fold change of each transition rate is the sum of each patient’s prognostic variables multiplied by the effects. Personalized outcome predictions are subsequently computed by integrating the time-dependent transition rates, multiplied by their predicted fold changes. d. Qualitative comparison of predicted 3yr survival versus observed outcome for all patients. Patients predicted to have high probability of survival typically have long observed survival times. This is quantified by the concordance C, indicating that the survival times of 72% of patients were correctly ranked by the model. Similarly, at three years after diagnosis only 28% of patients we e incorrectly predicted to be alive or dead. e. Mosaic plot of predicted 3-year survival across ELN categories. The size of each tile is proportional to the number of patients in each subgroup. f. Relative importance of risk factors for different transitions (as defined by arrows in part a as measured by the multistage model. A large proportion indicates that the variables of a given category are influential in determining the rate of the transition. The concordance C, shown as percentages across the top of the bar chart, indicates how accurately the model can predict each transition (50%=random, 100%=optimal).

### Personally tailored prognosis

In most Oncology practice currently, risk stratification is just that: grouping patients into broad strata based on a few predictor variables. In AML, the current prognostic standard is the European Leukemia Net (ELN) genetic scoring system ^11^, which defines four categories of disease risk based on 6 fusion genes, 3 point mutated genes and cytogenetic abnormalities. We find that individual risk in this AML cohort was instead continuous, with no obvious cut-points for stratification, suggesting that arbitrarily grouping patients of similar risk discards much prognostic information (Figure 2d). These survival estimates confirm the broad trends of known ELN risk groups; however a third of patients have survival predictions deviating more than 20% from the ELN stratum (Figure 2e).

From the multistage model, we can quantify how much each of the broad classes of predictor variables contributes to explaining population variance observed in the different possible endpoints of treatment (Figure 2e, Supplementary Table S2-6). We find that clinical and demographic factors, such as patient age, performance status and blood counts, exerted most influence on the rates of mortality in either stage. Genomic features, be they copy number changes, fusion genes or driver point mutations, most strongly influenced the dynamics of the disease remission and relapse.

These estimates represent the contributions of the various categories of predictors to outcomes of treatment *at the population level*. At the individual level, we can score each patient for his or her risk along these dimensions of predictor variables. What emerges is the considerable heterogeneity in personal risk profiles across the cohort (Figure S1). The heterogeneity of risk profiles and the variable impact they have on the different AML outcomes combine to generate a kaleidoscope of predictions for patients’ journeys through therapy (Figure 3). Thus, there are distinct groups of patients for whom we can confidently predict long-term survival in first remission, or death after relapse, or death without achieving remission, manifesting as swathes of purple, green or pink respectively in Figure 3. Reassuringly, these predictions square well with what actually happened to the patients (status lines and circles in Figure 3). It is these patients for whom the personally tailored predictions have much confidence. There are, however, some patients for whom there is genuine uncertainty about outcomes even with the full model. These patients have predicted survival curves that deviate little from the population average.

**Figure 3.**
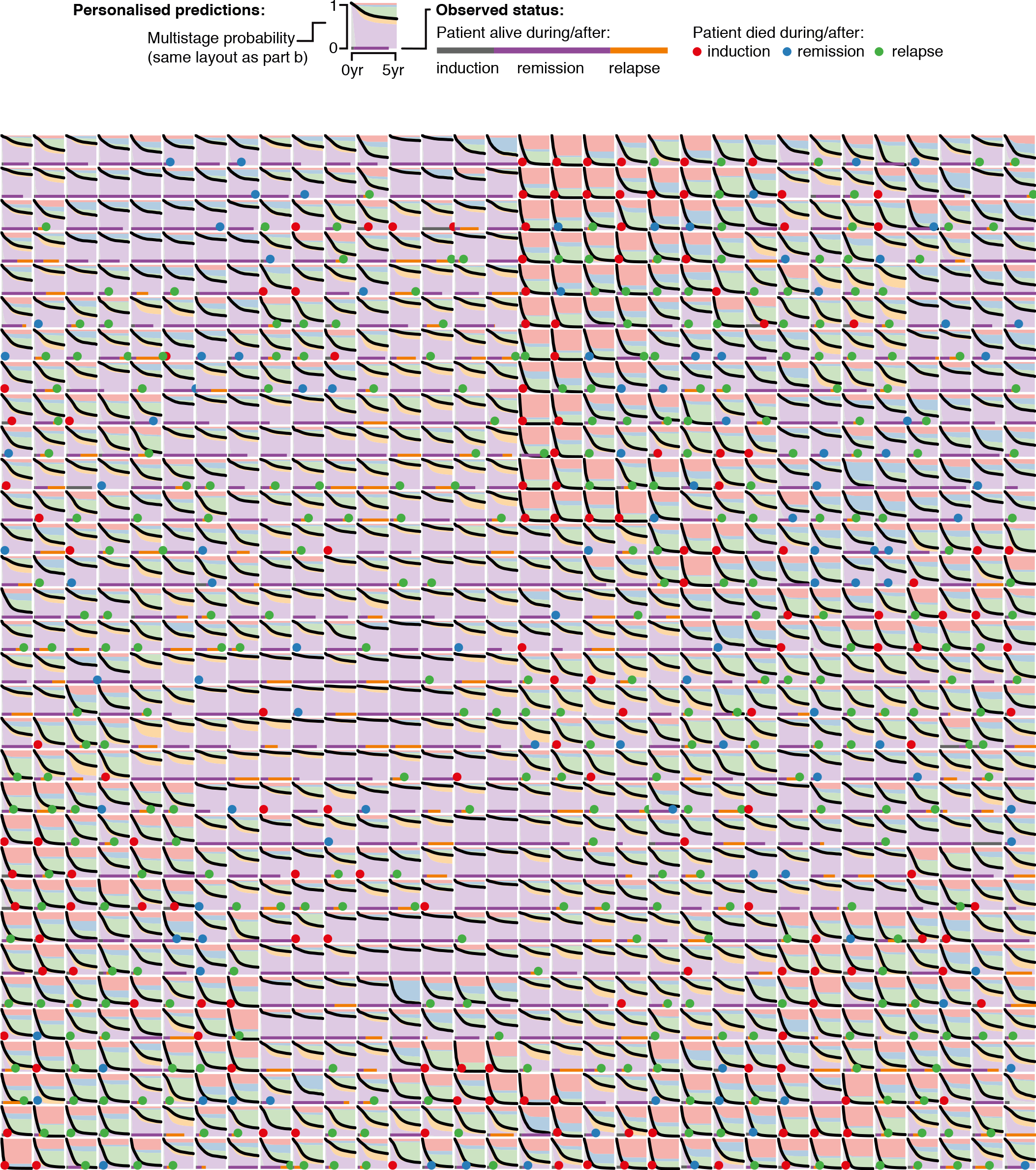
Multistage outcome predictions for 1024 patients. Cross-validated risk predictions and observed statuses for 1024 patients, arranged along a Hilbert curve. This has the property that patients with similar AML subtype and risk constellation are grouped together in the 2-dimensional space (compare Figure 2 for constellations of risk factors). For each individual patient, the survival curves predicted by the multistage model are shown, with the competing outcomes colored as in the legend and **Figure 1b**. What actually happened to the patient is shown as a line across the base of the graph, with a filled circle indicating the patient died, its color indicating the mode of death. Note that there are many patients for whom one color dominates the diagram, indicating that the probability that a particular event occurs is very high. Reassuringly, for such patients the observed outcomes are highly concordant with the cross-validated predictions and occur at frequencies matching the predicted probabilities.

### Personally tailored therapeutic decision support

Practically all cancer treatments are either invasive (surgery, radiation therapy, stem cell transplants) or have strong side effects (chemotherapy, targeted inhibitors and immunotherapy). Therefore an accurate assessment of a patient’s disease-associated risk, contrasted with the potential risks and benefits of each treatment option, is mandatory for evaluating different therapies. In AML, one of the therapeutic dilemmas is deciding which patients should be offered allogeneic hematopoietic cell transplants (allografts), and whether this should be in first complete remission or after relapse ^10,11^. With 10-20% transplant-associated mortality and high risk of long-term graft-versus-host disease, it is clear that allografts are reserved for high-risk patients, where disease-associated mortality exceeds treatment-associated mortality.

The above calculations have shown that a detailed survival analysis reclassifies a substantial fraction of patients into low and high risk (Figure 2e). This implies that potentially also the optimal treatment for these patients may differ. It also emerged that a patient’s risk represents an aggregation across multiple facets of the disease. Thus, two patients can both have an overall intermediate probability of death but arrive at this through different risk contributions: one might be older and more frail but have a leukemia with generally favorable genomic features; the other might be young and fit but with a leukemia carrying many adverse driver mutations. Intuitively, a clinician will favor the more intensive allogeneic transplant in the latter, fitter patient while preferring standard chemotherapy in the older patient at higher risk of treatment-related mortality.

A knowledge bank enables us to compute outcome under different treatment scenarios, thereby directly assessing risks and benefits on a per patient basis. In order to be able to accurately quantify the effect of a therapy, the treatment needs to be randomized. In this cohort 711/1540 patients received an allograft, however the decision to perform a transplant was only partially randomized^15^. Therefore the following calculations are only indicative and need to be further validated. However, the limited availability of a matched donor introduces a quasi-randomization, which at least allows for deriving meaningful effect sizes.

We illustrate these calculations using two patients from the cohort (Figure 2; other representative patients illustrated in Figure S2). The first was a 29-year woman with t(8;21) and no other driver mutations: favorable risk by ELN criteria ^11^. Under a strategy of chemotherapy in CR1 with salvage allograft after relapse, we predict her chance of 3-year survival to be 86% (CI_95%_: 78-91%) (Figure 4a). In contrast, with allograft in CR1, we estimate her overall cure rate to be 88% (79-93%) (Figure 4b), with the decrease in probability of relapse matched by the increase in non-relapse mortality with transplant. Hence there is no indication for an up-front allograft for this patient, with only 2 percentage points difference in predicted survival (CI_95%_: −3 to 7).

**Figure 4.**
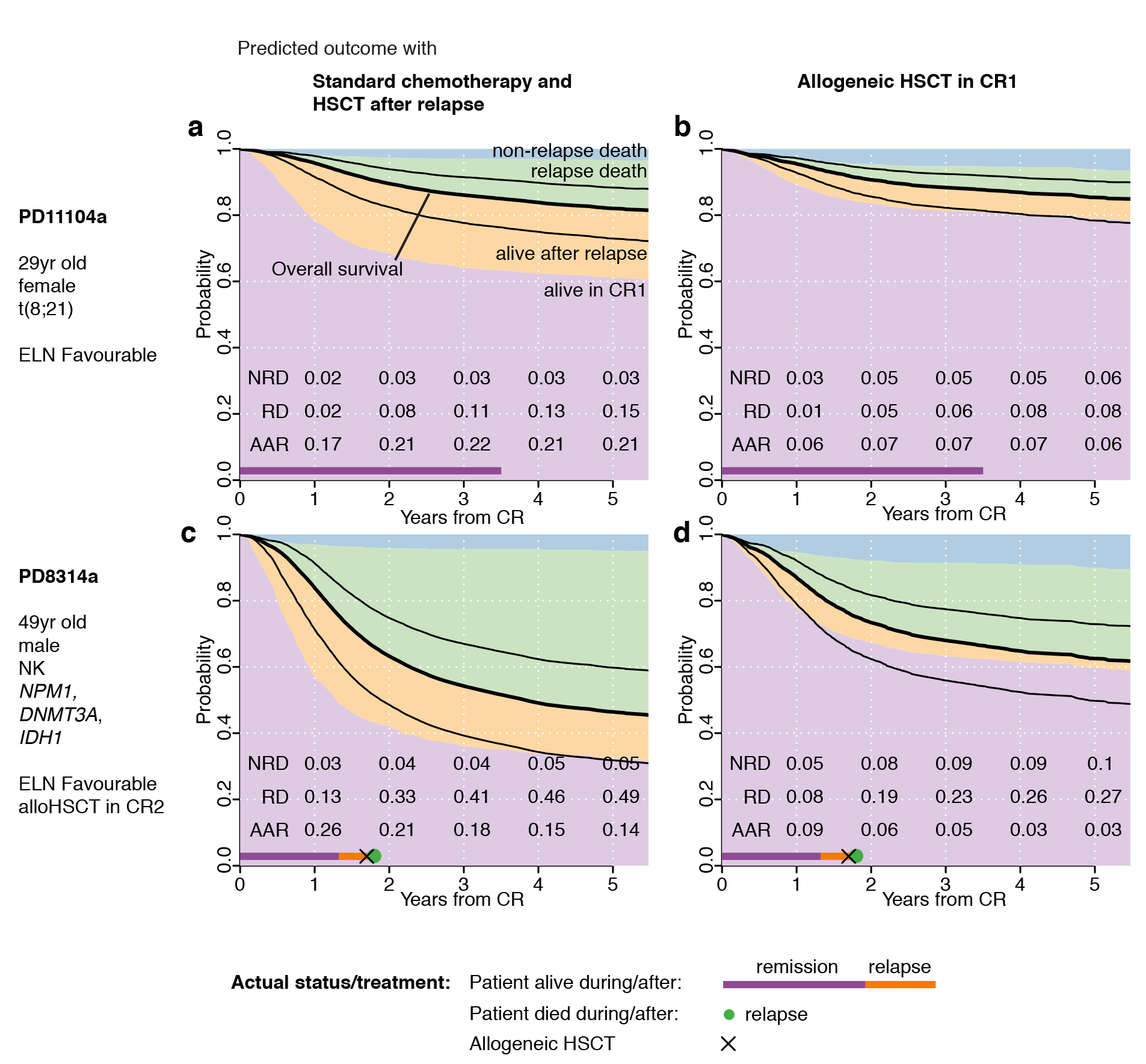
Individualized risk exemplified for 2 patients. a. Sediment plot showing predicted multistage probability after remission for patient PD11104a under a management strategy of standard chemotherapy in CR1 with intended allograft after relapse. Predictions shown are based on models where the given patients were excluded for training; the bar at the bottom denotes the observed outcome (as for **Figure 2**). The patient was alive at the last follow-up 3.5 years after achieving first complete remission. Numbers at the bottom indicate the probabilities of non-relapse death (NRD), post-relapse death (PRD) and being alive after relapse (AAR) at years 1 to 5 from achieving complete remission. b. Multistage probability for PD11104a in the scenario of an allograft in first complete remission. c. Same as a for patient PD8314a. The patient relapsed after 1.2 years and died soon after. d. Same as **b** for patient PD8314a.

The second patient was a 49-year old male with mutations in *NPM1, DNMT3A, IDH1* and normal karyotype. Under ELN criteria, his risk also classifies as favorable, and he would not currently be recommended for allograft in first CR. With standard chemotherapy as first-line therapy, we estimate his 3-year survival probability at 55% (41-67%), compared to 68% (55-77%) for allograft in CR1 (Figure 4c–d). Thus, his disease is not especially favorable risk when all predictive information is considered. Furthermore, the absolute risk reduction associated with an up-front allograft is estimated at 13 percentage points (3-24%). This is equivalent to curing 1 additional patient for every 7 (4-26) so treated, despite current guidelines recommending standard chemotherapy.

### Population-based gains from personally tailored decision support

We now extend the analysis for the two patients described above to all those under 60 years of age in the cohort to assess how big the potential benefit of an early allograft could be (Figure 5a). On average, we find that patients who are predicted to have poor prognosis, more than 60-70% chance of mortality at 3 years, are most likely to benefit from allogeneic transplantation in first remission, a finding captured in current clinical guidelines. However, while this may be valid at the population level, it does not necessarily translate to the level of the individual patient, since there is considerable spread of patient estimates around the population average.

Overall, we estimate that 12% (124/995) of patients in CR1 aged <60 years would gain more than 10 percentage points improvement in survival at 3 years with an allograft in CR1 compared to standard chemotherapy (number needed to treat <10; Figure 5b). These leave-one-out predictions agree with the observed differences in outcome (Figure 5c). Only 29 of these 124 patients are identified as adverse risk by current guidelines, with most being intermediate and some even favorable risk. Furthermore, 57% (302/534) patients classified as adverse or intermediate risk by ELN criteria, and therefore strongly considered for allograft in CR1 under current clinical guidelines ^11^, are predicted to derive <5 percentage points improvement in survival from up-front allografts. Clearly, then, a knowledge bank approach might change management in a large fraction of patients compared to current practice guidelines.

**Figure 5.**
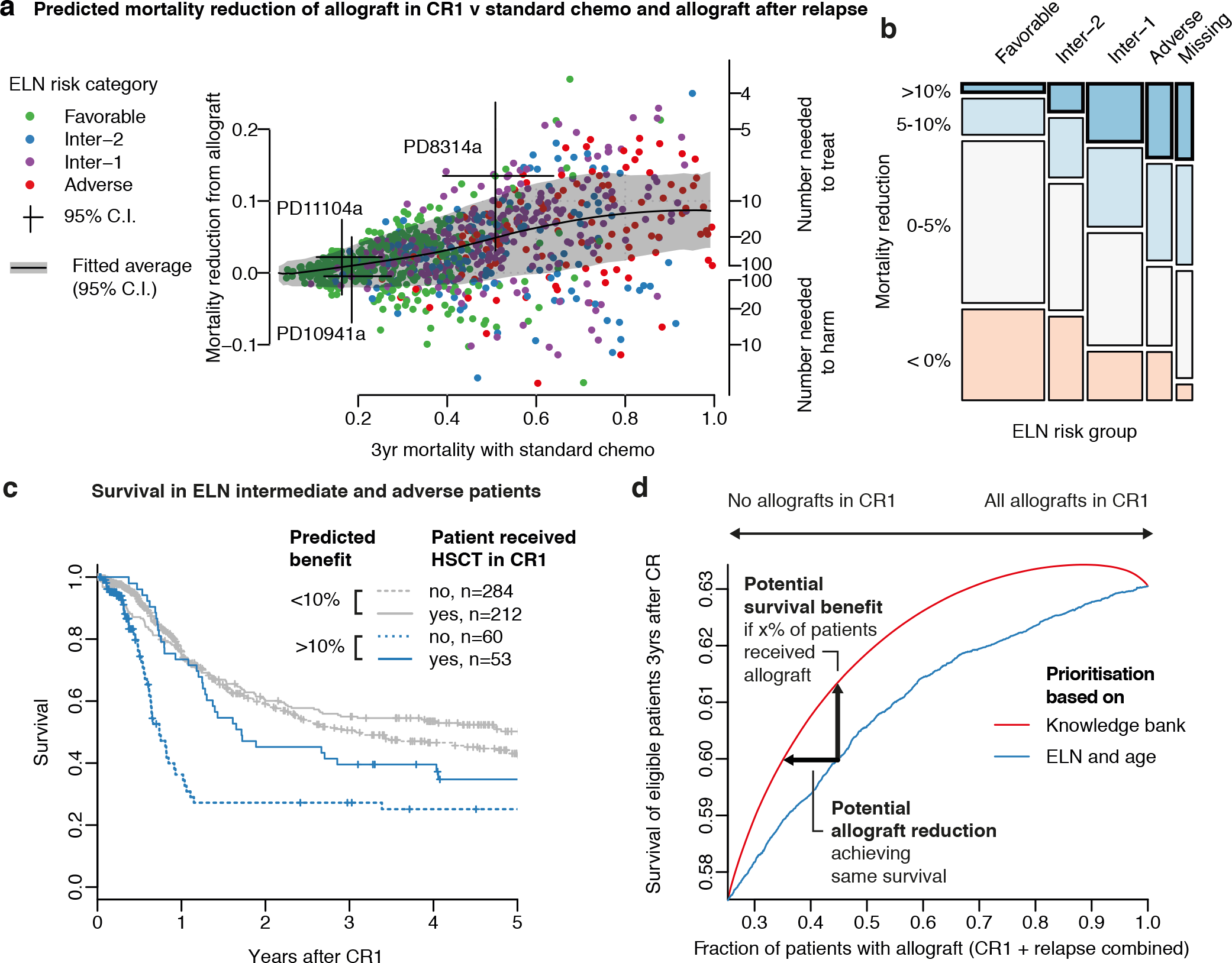
Benefit of allograft in CR1 vs after relapse. a. Predicted three-year absolute mortality reduction by allografts in CR1 over standard chemotherapy in CR1 and allograft after relapse (y-axis). This is shown in relation to three-year mortality without allograft as a measure of overall risk (x-axis). Points shown are for patients < 60yr of age in CR1 (n=1,109), who would generally be considered eligible for allogeneic bone marrow transplants. b. Mosaic plot of absolute survival benefit by an allograft in CR1 over standard chemotherapy and allograft after relapse at 3yrs after CR1 versus ELN risk category. 13% (146/1,109) patients have a predicted survival gain greater than 10% (‘high benefit’, bold lines). c. Kaplan-Meier curves for patients with high (>10%, light blue) and low (<10%, light red) predicted benefit of early allograft (cross-validated), each with and without allograft in CR1. Patients with favorable ELN risk showed a similar trend, but received fewer allografts, thereby skewing the results and were therefore excluded from this comparison. Data for all patients stratified by ELN risk group are shown in **Figure S3**. d. Predicted overall survival at 3yrs as a function of total number of allografts (in CR1 + after relapse). Patients are ranked left to right on the x axis from those most likely to benefit from transplant to those least likely to benefit, as judged by current guidelines (solid blue line), the current knowledge bank (solid red line), a knowledge bank of 10,000 patients (dashed red line) or a knowledge bank of optimal size (dotted red line). The total number f transplants performed is based on the assumption that half of relapsing patients treated with chemotherapy alone in CR1 manage to receive a post-relapse transplant (the fraction observed in this cohort). The distance between the curves shows that survival could be improved by a knowledge bank approach by ~1.3% using a fixed number of allografts, or that a similar survival could be achieved with ~10% fewer transplants.

The hypothetical population level benefit if treatment decisions were based on the above calculations can be quantified as either improvement in expected survival for a fixed number of allografts in CR1 or reduction in the number of allografts in CR1 needed to achieve the same overall survival (Figure 5d). In USA, ~30% of patients with AML receive an allograft ^18^ If the 30% to receive an allograft in CR1 were chosen using an optimal knowledge bank rather than current guidelines, we estimate survival rates across the cohort would increase ~1.3 percentage points (60% to 61.3%). While this seems small, it is to be compared with the average of 5% survival gain offered by a stem cell transplant in CR1.

Alternatively, to maintain the same survival as achieved by current guidelines, ~10% of patients could be saved from an allograft if allocated using an optimal knowledge bank. Under current guidelines, 44% young patients would receive a transplant, broken down as 30% in CR1 plus 14% post-relapse. In contrast, using a knowledge bank approach, 35% patients overall would receive a transplant, as 16% in CR1 plus 19% post-relapse. Similar overall gains from a knowledge bank approach were found across a range of assumptions for risks and benefits of transplant (Figure S4).

### Portals for exploring decision support predictions

The preceding sections demonstrate that the complex and multifactorial interrelationships among genomic variables, clinical predictors and cancer outcomes can be learnt with a sufficiently comprehensive knowledge bank. Since the underlying survival models are complex, diagnostic laboratories may need to provide personalized portals into a given patient’s cancer genome.

Our dataset is not appropriate for direct clinical use, as the algorithm has not yet been prospectively validated and sequencing was performed using a research pipeline. Nonetheless, as a research tool, we have created a portal within our website ^19^ (http://cancer.sanger.ac.uk/aml-multistage; user: reviewer, password: Amlpred42) that allows outcome predictions to be generated based on this dataset for user-defined constellations of genomic features, clinical variables and treatment strategies (Figure S5). The underlying algorithm is capable of imputing missing variables and computes confidence intervals for each prediction.

### The knowledge bank

We explored how both the breadth of genomic profiling and the sample size of the knowledge bank impact on the accuracy of outcome predictions for individual patients. The explained risk grows linearly with the *average number of driver mutations present in each patient* (Figure S6a), a relationship underpinned by theoretical arguments (Supplementary Methods S5.3.2). Some genes, by virtue of their frequency and/or the magnitude of their prognostic effect, are more informative than others. We have ranked AML genes by their predictive utility (Figure S6b), such that laboratories with limited sequencing capacity can choose the most useful subset of genes to test. The effects of missing mutation data on confidence intervals of patient prediction can be explored in the web portal.

The other critical factor to accurate risk profiling is the sample size of the knowledge bank. Using subsampling analyses and simulations from the AML data, we found that prognostic accuracy steadily increases with larger sample sizes, albeit following a law of diminishing returns (Figure 6a). As a rule of thumb, to detect a moderate-sized prognostic effect of a given cancer gene, say an increase of 50% in relative risk, the knowledge bank needs ~50-100 patients with that mutation (Figure 6b, Figure S7a). Thus, for a gene mutated at 10% of patients, a training set of 500-1000 patients would suffice, but for a 1% gene, a cohort of 5,000-10,000 would be needed. These simulations match theoretical expectations ^20,21^ (Supplementary Methods S5.4.2; Figure S7b).

**Figure 6.**
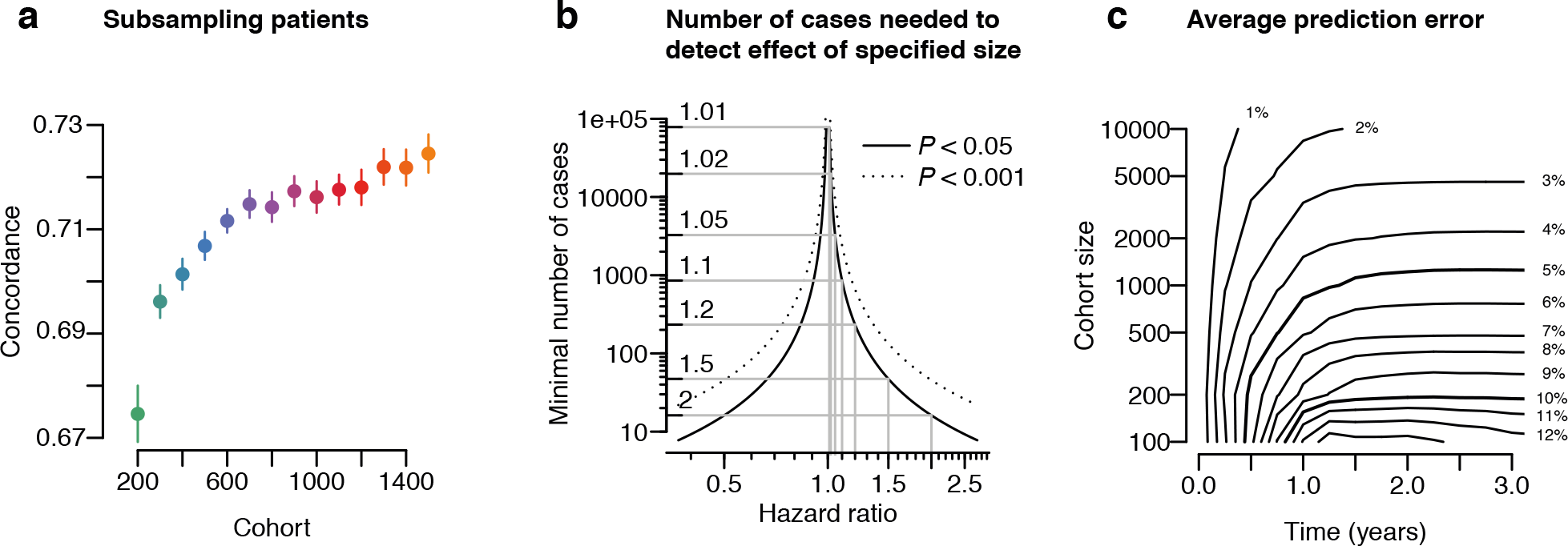
Extrapolations and power calculations. a. Subsampling of the number of patients reveals a steady, but saturating increase in prognostic accuracy. b. Graph relating the effect size (hazard ratio) of a prognostic variable to the absolute number of patients with the given factor required to reach significance (solid line: P < 0.05; dotted P < 0.001). c. Average prediction error as a function of survival time (x-axis) and training cohort size (y-axis).

The standard error of our survival predictions 3 years after CR is about 6%. When using predictions for supporting therapeutic decisions about a specific patient, this uncertainty limits the ability to confidently discriminate small differences in survival. With 1,000 cases, we could achieve an average absolute prediction error for an individual patient of approximately 5 percentage points, which could be brought down to 2 percentage points with 10,000 cases (Figure 6c).

## DISCUSSION

Here, we have evaluated the promise of precision medicine, building statistical models that can generate personally tailored clinical decision support from all available prognostic information in a knowledge bank. From a database of 1540 patients, we can make considerably more informative and more accurate statements about an individual’s likely journey through AML therapy than the current standards in clinical practice. Our approach enables us to compare the likelihood of favorable outcomes under different treatment scenarios, providing information that can support genuinely personalized decision-making. While we have focused on AML in this analysis we believe that the same logic applies to knowledge banks from other cancer types, which will be generated as genomics enters healthcare and healthcare becomes digitized.

Building and maintaining clinical-genomic knowledge banks is a formidable challenge, especially for solid tumors where the genome can be considerably more complex than AML. Initially, knowledge banks could be seeded from clinical trials cohorts, as we did here, since these will have high quality clinical data and state-of-the-art therapies. Our power calculations suggest, however, that most clinical trials would not be powered to detect gene-drug interactions involving genes mutated in <20% patients. Additionally, knowledge banks will need to include patients who are representative of the wider cancer population to enable meaningful extrapolation to real-world clinical practice. We believe, therefore, that data from post-marketing surveillance of newly approved therapies and from hospital-based practice would further enrich knowledge banks. This would require investment, though, to ensure that clinical and genomic data were accredited and sufficiently robust to underpin decision support for individual patients. Finally, statistical methods will need to evolve for making personalized predictions from complex datasets, requiring community collaboration and competition ^22^.

Whether the returns justify this investment will be contentious. Here, we have illustrated how the calculations play out for a specific clinical conundrum. If 30% of intensively treated AML patients were allografted in CR1, we estimate that 3-year survival could be increased by 1.3 percentage points. We should not be surprised at how modest the gain is – for the bulk of patients, we predict only small improvements in survival with early allografts (Figure 5b). What may be more important is the more accurate allocation of a precious resource, since we can potentially save 10% of patients from an allograft while maintaining the same overall survival as for current guidelines. Not only would this reduce morbidity from chronic graft-versus-host disease, at US$100,000-200,000 per allograft ^23^, the potential monetary savings would far outstrip the costs of the genomic screens.

There is a tension between maintaining the precepts of *evidence-based medicine* while sharpening the focus on the individual with *precision medicine* ^24^. Here, we have demonstrated how knowledge banks can resolve this tension, using the evidence base of thousands of patients to inform outcomes for the individual. The therapeutic choice we exemplified is binary: transplant versus chemotherapy in AML. The success of FLT3 inhibitors ^25^ potentially squares the number of available treatment options, and other novel agents will add further complexity. Knowledge banks could be a useful tool for clinicians navigating this complexity, but must remain evergreen as the therapeutic armamentarium expands and as our molecular understanding of cancer deepens. The logistic and regulatory hurdles, the scale needed and the costs of such an undertaking are daunting but not insurmountable.

## ONLINE METHODS

We performed targeted gene sequencing of 111 myeloid cancer genes ^17,26–28^ on DNA from leukemic cells in a cohort of 1,540 adults with AML who were treated with intensive therapies in three clinical trials run by the German-Austrian AML Study Group ^29–31^. The sample cohort, sequencing protocol, mutation-calling algorithms and definition of driver mutations are described in Ref. 16. In AML-HD98A, patients aged 18-61 years received induction chemotherapy with idarubicin, cytarabine and etoposide (ICE), followed by allogeneic transplants for intermediate-risk patients with matched related donor and high-risk patients; intensive consolidation chemotherapy for the remainder. Treatments were similar in AMLSG-07-04 but included randomization for all-*trans* retinoic acid therapy (ATRA) or not in induction. In AML-HD98B, patients ≥61 years received ICE±ATRA, with further therapy dictated by response. Median follow-up was 5.94 years.

We explored a range of statistical methods to build models of overall survival ^32,33^ including random survival forest regression; stepwise Cox proportional hazards model selection with either AIC or BIC penalty; complementary pairs stability selection based on LASSO penalized Cox proportional hazards models; random effects models with Gaussian random effects/ridge penalties; and random effects multistage models (Supplementary Methods S2-4). We found little prognostic significance to whether mutations were subclonal or clonal (Figure S8), and therefore do not consider this information in the multivariate models. All predictions shown are based on a leave-one-out basis; it is therefore informative to compare each prediction with the observed outcome in a given patient.

All predictions for individual patients reported here were made using models excluding that patient. To maximize reproducibility, details of statistical methods and all analysis code used are provided in Supplementary Methods and as a git repository (http://www.github.com/mg14/AML-multistage).

## ACKNOWLEDGEMENTS

We thank Chris Holmes for stimulating discussions. This work was supported by the Wellcome Trust (077012/Z/05/Z). PJC has a Wellcome Trust Senior Clinical Research Fellowship (WT088340MA). EP is supported by an EHA early career fellowship. Supported in part by grants 01GI9981 and 01KG0605 from the German Bundesministerium für Bildung und Forschung (BMBF), grant 109675 from the Deutsche Krebshilfe, and by project B3 and B4, Sonderforschungsbereich (SFB) 1074 funded the Deutsche Forschungsgemeinschaft; HD is coordinating investigator of SFB 1074; LB is a Heisenberg Professor of the DFG (BU 1339/3-1). We gratefully acknowledge Daniela Weber for clinical data managing, Veronica Teleanu for assistance in cytogenetics data classification, and Dr. Sabine Kayser for assistance in morphologic evaluation. We are grateful to all members of the German-Austrian AML Study Group (AMLSG) for their participation in this study and providing patient samples; a list of participating institutions and investigators appears in the Appendix to the companion paper. AMLSG treatment trials were in part supported by Amgen and DKH Grant: 109675.

### Author contributions

MG developed all methods, analyzed data and wrote the manuscript and supporting information, with input from EP and PJC. EP prepared and curated genetic and clinical data. IM analyzed TCGA data. RS, HD, KD, LB, VIG, PP, MH, FT, AG alongside all contributing institutions to the study group (AMLSG) recruited patients in this study, collated and contributed clinical data. NB, PG and UM provided input into analyses and interpretation of results. EP, KD, HD, RFS and PJC initiated the study. PJC and HD wrote the manuscript, and are joint corresponding authors.

